# Balanced Oscillatory Coupling Improves Information Flow

**DOI:** 10.1101/030304

**Authors:** Erik J. Peterson, Bradley Voytek

## Abstract

All animals are able to rapidly change their behavior. The neural basis of such flexibility requires that groups of distant neural ensembles rapidly alter communications with selectivity and fidelity. Low frequency oscillations are a strong candidate for how neurons coordinate communication via the dynamic instantiation of functional networks. These dynamic networks are argued to rapidly guide the flow of information, with the presumption that stronger oscillations more strongly influence information flow. Surprisingly, there is scant evidence or theoretical support for how oscillatory activity might enhance information flow. Here we introduce a novel computational model for oscillatory neural communication and show that, rather than the strength of the oscillation, it is the balance between excitatory and inhibitory neuronal activity that has the largest effect on information flow. When coupling between an oscillation and spiking has balanced excitatory-inhibitory inputs, information flow is enhanced via improved discriminability between signal and noise. In contrast, when coupling is unbalanced, driven either by excessive excitation or inhibition, information flow is obstructed, regardless of the strength of the oscillation. A multitude of neuropathologies, including Parkinson’s disease, schizophrenia, and autism, are associated with oscillatory disruptions and excitation-inhibition imbalances. Our results show that understanding the distinction between balanced and unbalanced oscillatory coupling offers a unifying mechanistic framework for understanding effective neural communication and its disruption in neuropathology.

## Introduction

In a loud, crowded room, determining who is saying what to whom and when can be a daunting task. This spoken language communication problem pales in comparison to the communication problem between neural ensembles in the brain. Brains have countless possible overlapping anatomical networks of varying sizes and strengths. One dominant theory for how neurons dynamically route information within and between these networks with fidelity, depending on behavioral demands, relies on neural oscillations [Fries, 2005].

Neural oscillations are self-organized phenomenon that play an important role in cognition and neural communication [Buzsáki, 2006]. They track perception [Spaak et al., 2014], attention [Engel et al., 2001, Fries et al., 2001, Saalmann et al., 2012, Szczepanski et al., 2014], learning [Schaefer et al., 2006, Litwin-Kumar and Doiron, 2014, Ainsworth et al., 2012, Tort et al., 2009], memory [Jensen et al., 1996, Fell and Axmacher, 2011, Voytek and Knight, 2010, Voytek et al., 2013, Lisman and Jensen, 2013], and cognitive control [Cooper et al., 2015, Voytek et al., 2015], among many other behaviors and cognitive states [Wang, 2010, Womelsdorf et al., 2007a, Buschman et al., 2012]. Mechanistically, oscillations bias neuronal spiking [Womelsdorf et al., 2007b, Jia et al., 2013, Sohal et al., 2009] and are thought to aid in information flow between brain regions [Siegel et al., 2015, Buehlmann and Deco, 2010, von Nicolai et al., 2014, Voytek and Knight, 2015]. Furthermore, they have been implicated in nearly every major neurological and psychiatric disorder [Herrmann and Demiralp, 2005, Uhlhaas and Singer, 2006, Voytek and Knight, 2015, de Hemptinne et al., 2015, Allen et al., 2011, Khan et al., 2013].

The nature of excitatory (E) and inhibitory (I) neuronal interactions induce oscillations naturally and spontaneously [Buzsáki, 2006, Wang, 2010]. Balanced excitatory (E) and inhibitory (I) activity have been shown to be crucial components in the studies of effective neural communication [Vreeswijk and Sompolinsky, 1998, Ostojic, 2014], assembly formation [Litwin-Kumar and Doiron, 2012], working memory [Lim and Goldman, 2013], and neural computation [Murphy and Miller, 2009, Litwin-Kumar et al., 2011, Abbott and Chance, 2005]. By ‘balanced E-I interactions’ we mean that increases in excitatory firing are rapidly and exactly countered by equivalent increases in inhibition [Atallah and Scanziani, 2009], leaving the network in a fluctuation driven state [Renart et al., 2007].

While oscillations have been shown to have a facilitative role in cognition; they are also known to obstruct behavioral functioning [de Hemptinne et al., 2015]. We argue that both their facilitative and obstructive roles can be understood as a consequence of their biophysical origin. In order to understand the functional *and* pathological role that oscillations play in information flow, we need to first create a simple, generalizeable neural framework for linking oscillations and information.

We begin with a form of oscillatory entrainment or modulation of neural activity known as phase-amplitude coupling (PAC). Generally, PAC is thought to reflect rhythmic changes in neural excitability [Canolty and Knight, 2010]. When phase modulates firing rate directly, as opposed to entraining another oscillator, PAC becomes analogous to spike-field coupling and is often known as “high-gamma PAC”. PAC between theta oscillations (6-12 Hz) and the high gamma frequency range (80-300 Hz) has been argued to enhance neural information flow [Siegel et al., 2015, Buehlmann and Deco, 2010, von Nicolai et al., 2014, Voytek and Knight, 2015], facilitate long-term potentiation [Hölscher et al., 1997], and improve behavioral performance [Tort et al., 2009]. Seemingly paradoxically, the presence of PAC has also been linked to clinical outcomes such as Parkinson’s disease [de Hemptinne et al., 2013], schizophrenia [Allen et al., 2011], and autism [Khan et al., 2013], with clinical deep brain stimulation in Parkinson’s disease associated with a reduction in Parkinsonian symptoms and a concomitant reduction in (presumably) pathological PAC [de Hemptinne et al., 2015].

In this simulation study, we model PAC as periodic modulation of an asynchronous, rate-based, population code. Simplified rate models of Poissonic firing have proved to be pivotal in isolating key principles of neural function [Heeger et al., 1996, Reynolds and Heeger, 2009, Mazurek and Shadlen, 2002, Bays, 2014]. This simplified approach also allows our model to encapsulate a broad class of ways in which excitatory or inhibitory oscillations might modulate firing rates [Silver, 2010]. We then use information theory to separate models that facilitate communication from those that obstruct it. We find that only models based on multiplicative gain control–a class of models that implicitly require strong and balanced E-I inputs [Chance et al., 2002, Vreeswijk and Sompolinsky, 1998, Brunel, 2000, Abbott and Chance, 2005, Womelsdorf et al., 2014]–facilitate information flow. Importantly, we identify a plausible mechanism by which this information enhancement occurs, whereby E-I balanced oscillations improve the ability of downstream neural populations to discriminate between input signal and noise.

## Methods

### Neural architecture

Our formalization of PAC is well characterized by an expanded version of the E-I driver-modulator framework (see [Abbott and Chance, 2005]). Our driver population is direct excitatory stimulation (population *s*, (Figure 1A)), which exhibits asynchronous naturalistic firing patterns (explained below). Modulation takes the form of slow (6 Hz) endogenous oscillatory activity. While previous models used balanced E-I input to implement gain control [Abbott and Chance, 2005]–*i.e*., multiplicative scaling (modulation) of *s*-our model also considers modulation by strong excitation and strong inhibition (Figure 1B). Including all three forms of modulation corresponds to a complete implementation of “neural arithmetic”, a well-established, perhaps canonical, view of neural computation [Silver, 2010]. In this view, changes to excitatory conductance (generically labeled with a *g* in the equations below) leads to a rightward shift of the neurons’ tuning curves, akin to “addition” (Figure 1B, middle). Increases in inhibitory conductance lead to a leftward shift synonymous with neural “subtraction” (Figure 1B, bottom). Finally we simulate gain control as a multiplicative process characterized by a leftward and upward warping of each neurons’ response function (Figure 1B, top).

**Figure 1:**
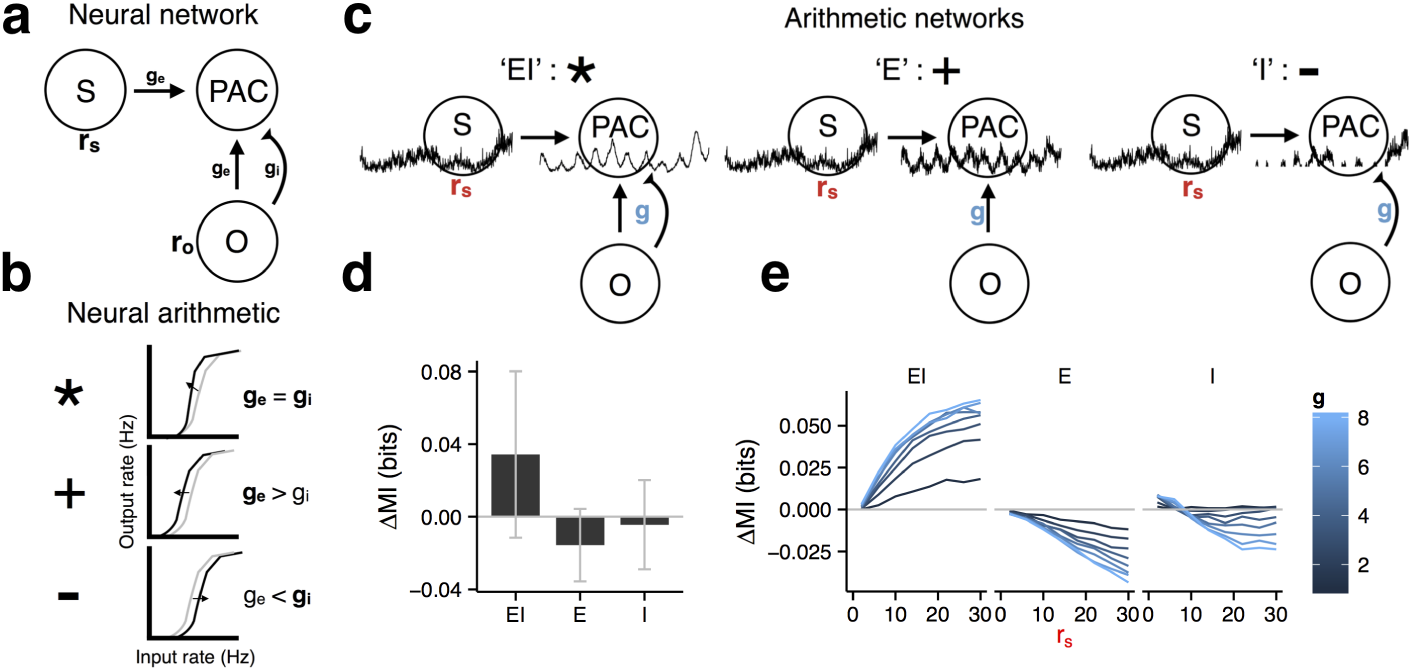
Phase-amplitude coupling (PAC), excitatory-inhibitory interactions, and their mutual effect on information flow in a simplified neural circuit. **a)** Diagram of a simple neural model capable of generating PAC. This model contains an excitatory stimulus population (*s*), which functions as a strong driver of the downstream populatio, as well as a modulatory oscillatory population (*o*), both of which synapse onto a third target population, *PAC*. All connections are completely characterized by a synaptic strength of *g*, and the population firing rate is given by *r*. **b)** Diagram of “neural arithmetic”, characterized as a single neuron’s input-output firing rate curve (*i.e*., *F-I* curve). The top panel depicts balanced increases to both excitation and inhibition, which lead to a multiplicative increase in firing, *i.e.,* gain control (*ge* = *gi*). Increasing only excitation (*g_e_* > *g_i_*) leads to leftward shift of the input-output firing curve and is synonymous with addition (middle). Increasing only inhibition (*g_i_* > *g_e_*) is equivalent to a rightward shift, or subtraction (bottom). **c)** Illustration of each PAC in “arithmetic network” form. Each model is an implementation of a subset of the full model outlined in **(a)** but replacing traditional synaptic dynamics with linear changes to firing rate based on neural arithmetic operations – multiplicative (EI), addition (E), subtraction (I). Example simulated LFP time courses, of both s and PAC, are shown. **d)** Overall change in mutual information (AMI) for all three PAC models (see Methods). Error bars are standard deviations. **e)** Change in mutual information (∆MI) plotted as a function of stimulus firing rate *r_s_* and synaptic strength *g* (blue).

Spiking in all neural populations is treated statistically, where each neuron acts as an independent time varying, *i.e.,* non-homogeneous, Poisson process. Each population has 100 neurons, though explored values between 50 and 500 do not alter the qualitative results. Beyond 500 neurons the results do change, and the advantage of gain to information flow rapidly declines, and begins to become obstructive. This size dependence suggests PAC is effective when applied only to small assemblies of neurons. Experimentally, it has been shown that assemblies in the 100-500 size range form the basis of neural computation [MacLeod and Laurent, 1996, Litwin-Kumar and Doiron, 2014, Buzsáki, 2010].

The basic Poisson rate model is given by 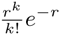 where *r* is the firing rate (traditionally designated *λ*) and *k* is the number of spikes per time step (1 ms for all models). Diffusion processes have been suggested as a reasonable approximation to the “naturalistic” firing patterns observed in early visual areas during passive movie viewing [Mazzoni et al., 2011, Barbieri et al., 2014]. As such, firing in the stimulus population was given by, 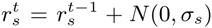 where *N*(0, *σ_β_*) is the normal distribution with *σ_s_* = 0.01*r_s_* and constrained so *r_s_ ≥* 0. Oscillatory firing was sinusoidal, where, 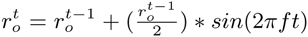 where f is 6 Hz. *r_o_* was fixed at 2 Hz. To constrain the firing models closer to biological reality, an absolute refractory period of 2 ms was enforced. Note that while rate terms are explicitly functions of time (*i.e., r*(*t*)), we use the superscript notation *r^t^*.

The oscillation rate 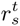 modulates the stimulus 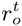 with coupling strength *g*. To better simplify the initial presentation, the overall model is depicted with independent E and I modulation terms, *g_e_* and *g_i_* (Figure 1A). However *g_e_* and *g_i_* can be replaced with a single term *g.* In the gain condition, *g_e_* = *g_i_.* In the excitatory and inhibitory conditions each term is varied independently but over the same range (Figure 1A). This new common *g* term then defines the coupling strength for all models (see below). Values of *g* were bound between 1-8, consistent with previous modeling efforts [Brunel, 2000].

Our simplified “arithmetic networks” (Figure 1) simulate changes to excitatory or inhibitory conductance by direct linear manipulation of the PAC population’s rate, *r_m_*. In *EI*, gain modulation was defined by 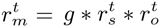 whereas *E* and *I* modulation were defined by 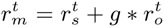 and 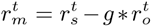 respectively (see Figure 1A for a graphical depiction). Results in the figures are averages of 100 random simulation runs, each defined with its own unique but randomly initialized stimulus pattern and Poissonic neural realizations.

### Information estimation

Information content was estimated from the *PAC* and stimulus s populations using summed population activity at every time-step (*i.e*., in 1 ms intervals). To allow for reliable calculation of the conditional probabilities necessary for information theoretic calculations, the summed rates were discretized into 8 integer levels. Integer levels between 4 and 30 were considered, but did not alter the patterns of results we report here. Using this new activity “alphabet”, entropy and mutual information (MI) were subsequently calculated with the pyentropy library [Ince et al., 2009], using the Panzeri–Treves method [Panzeri and Treves, 1996] of correcting for downward bias in estimating entropy *H* introduced by finite sampling of each time window.

### The definition of facilitative and obstructive PAC

In line with the conceptualization of PAC as enhancing neural communications, we offer a definition of facilitative and obstructive PAC based on a set of information theoretic inequalities. The precept is simple. Beneficial PAC should improve information flow while obstructive PAC should hinder flow. We then use the level of (Poissonic) noise present in the stimulus s itself as the baseline to compare against. We estimate this baseline by calculating the mutual information of two Poisson instantiations of the same inhomogeneous stimulus rate process, giving two spike trains *s* and *s′* (for more on the origin of *s* see above). Facilitative PAC, by definition, has higher MI than the baseline reference, *i.e*., Eq 1. In contrast, obstructive PAC is defined to have decreased information flow compared the baseline 2.

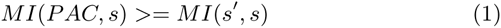

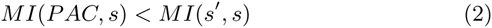

PAC-mediated changes to signal strength were assessed using d-prime 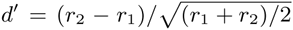 where *r_1_* and *r_2_* represent firing rates. As out underlying statistical model is Poisson the divisor in our *d*′ estimate replace uses average rate in place of the traditional variance.

### PAC estimation

The degree of phase-amplitude coupling in our models was assessed using the direct PAC estimator (see [Özkurt and Schnitzler, 2011]), a normalized version of the common modulation index [Canolty et al., 2006]. In brief, this method uses the summed product of instantaneous estimates of phase and amplitude to estimate degree of coupling. Instantaneous estimates are provided by the Hilbert transform [N. E. Huang et al., 1998]. All spectral estimation used simulated local field potentials (LFP; next section).

### LFP simulation

Simulated LFPs were constructed by convolving the population firing rate summed at each time-step with a double exponential kernel tuned to match fast AMPA channel kinetics. Changes to firing rate along with an exponential post-synaptic are a reasonable, albeit simple, approximation of an LFP recording [Buzsáki et al., 2012, Barbieri et al., 2014]. Simulated LFPs were used only in spectral analyses (Figure 2).

**Figure 2:**
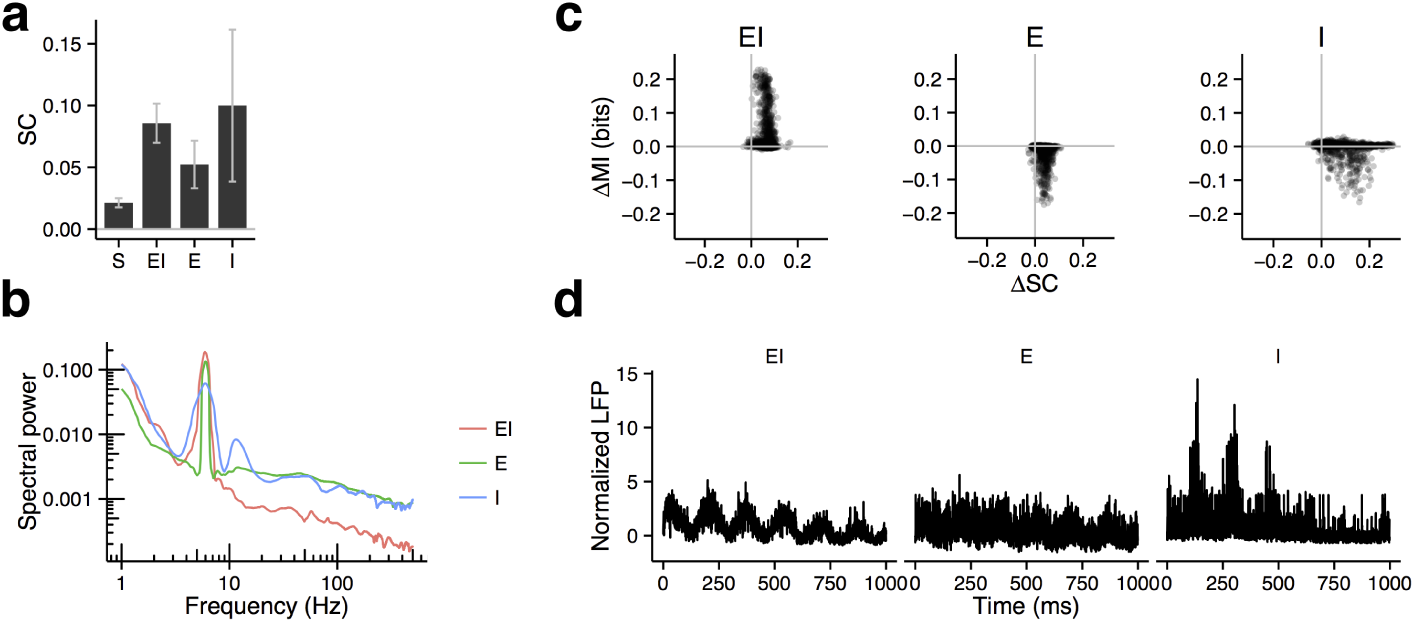
Spectral assessment of phase-amplitude coupling, matching the approach unused in past empirical studies, and its relation to mutual information flow. **a)** Average spectral coupling (SC) for all three PAC populations (*EI, E*, and *I*), as well as the stimulus population (*S*). Error bars are standard deviations. **b)** Change in PAC, ΔSC, (comparing PAC population to stimulus population) compared to the change in mutual information, ΔΜΙ, for all simulations and parameters (*N_total_* = 6400). Despite each condition having noticeable PAC as measured by SC **(a)**, and SC scaling with model parameters (see text), there was no significant correlation between PAC change and information flow change. **c)** Power spectrum of simulated LFPs from all three PAC models (*g* = 4, *r_s_* = 14) **d)** Simulated LFPs from **(c)**.

## Results

### Information and oscillatory modulation

Because E-I balance has been shown to be critical for effective neural communication, considering phase-amplitude coupling under a balanced E-I framework, specifically as a gain control process [Murphy and Miller, 2009, Abbott and Chance, 2005], would therefore be a fairly natural extension of past work. However empirical and simulation studies suggest populations of only excitatory [Cellulaire, 1992, Hansel et al., 1995] or only inhibitory neurons [MacLeod and Laurent, 1996, Wang and Buzsaki, 1996], as well as populations of both excitatory and inhibitory ensembles [Brunel and Wang, 2003, Geisler et al., 2005, Brunel, 2000, Börgers and Kopell, 2003, Tiesinga and Sejnowski, 2009], all can sustain robust oscillations across a range of frequencies (for a review see [Wang, 2010]). This suggests the oscillatory drive in PAC may come from E or I cells (middle, right panels in Figure 1a.), or from balanced E-I interaction synonymous with multiplicative gain (left panel in Figure 1a.). Therefore, we report here how each kind of entrainment, *E*, or *I*, or *EI*, altered information flow as function of entrainment degree (*g*, see *Methods*) and stimulus strength (*r_s_*).

Balanced inputs (*EI*) led to an increase in information flow (see ∆*MI* in *EI*, Figure 1**d**). In contrast, information flow declined when PAC was based on excitatory firing, modeled as a linear additive increase in rate, or as an inhibitory process, *i.e*., a subtractive decrease (see *E* and *I*, Figure 1**d**). Both the strength of coupling (*g*, blue) and the firing rate of the stimulus population (*r_s_*) monotonically enhanced each model’s effect on information flow. As *g* and *r_s_* increased in the *EI* model, so too did information flow. Again, in contrast *E* and *I* saw increasing declines (Figure 1**e**).

### Spectral measurement and mutual information

Phase-amplitude coupling between slow theta and high gamma in humans was first identified in recordings of cortical tissue [Canolty et al., 2006], and continues to be assessed with a range of largely equivalent spectral methods [Tort et al., 2010]. We applied one of these methods, the direct PAC estimator (see *Methods* and [Özkurt and Schnitzler, 2011]), to an LFP estimate derived from our simulated neural population. Partially confirming the validity of our model definition, we find that all models had a degree of spectral PAC (referred to as “SC” for spectral coupling PAC) above that present in the stimulus alone (compare all models to *S* in Figure 2**a**).

In the *E* and *I* models, SC positively correlates with degree of coupling *g* (*ρ_Ε_* = 0.32, *ρ_Ι_* = 0.51, *p* < 2 × 10^−16^). Interestingly, the strongest spectral PAC was driven by inhibition (*I* in Figure 2**a**), which has important implications regarding the nature and interpretations of experimentally measured PAC. In contrast, for the *EI* model there is a negative correlation between *g* and SC (*ρ_ΕΙ_* = −0.39, *p* < 2 × 10^−16^). These results suggest that under *EI* balance, increasing conductance actually reduces PAC. Notably, in all models the firing rate *r_s_* negatively correlates with SC (*ρ_EI_* = −0.48, *ρ_Ε_* = −0.60, *ρ_I_* = −0.50, with *p* < 2 × 10^−16^ all models). This relation is to be expected. The stronger the stimulus–the larger its firing rate–the the more oscillatory modulation is needed to effect it.

Changes in PAC, however, does not predict changes to information flow in any model (Figure 2**b**). Binning data by coupling strength *g* or firing rate *r_s_* does not alter this null result (not shown). This is somewhat surprising given that increased PAC is often associated with improved behavioral outcomes, except for pathological cases. This result hints at a more complex relationship between PAC and information flow. Spectral PAC methods, which relate band-passed regions of the power spectrum, may be insensitive to the spectral and temporal changes that accompany changes to information flow. SC measures were based on simulated LFPs (see *Methods*).

### Information flow in temporal and spectral domains

We showed (above) that changes to information flow, changes to PAC in our models, and changes to spectral coupling are model dependent. Specifically, we find that an increase in spectral coupling is not indicative of an increase in information flow (Figure 3). In fact, the opposite pattern was frequently observed (compare Figure 2 and 3). Notably, however, it is observed from the full spectral and temporal series that facilitative PAC has distinct temporal and spectral features compared to both obstructive models and the original stimulus population (Figure 2**c** and **d**).

**Figure 3:**
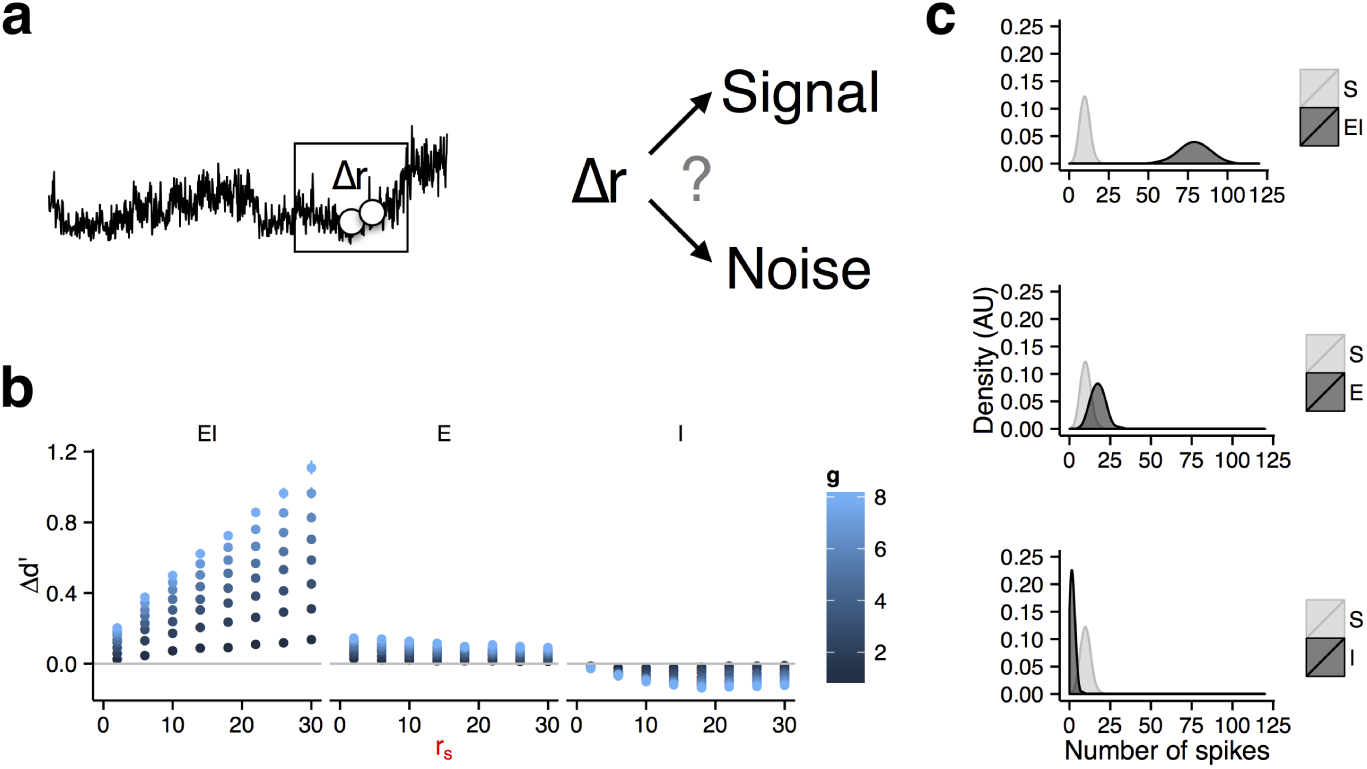
Signal detection theory explains the mutual information changes in our three models of phase-amplitude coupling. **a)** Illustration of rate coding as a signal detection problem. Under the rate-coding model, a critical aspect of decoding a rate-driven population is whether an instantaneous rate change is driven by the input signal, or noise fluctuations. **b)** In a rate-coding regime, we can assess the distinction between signal and noise by comparing moment-to-moment population firing rates using signal detection theory, using *d*′. We compare *d*′ values between the stimulus population, *S*, and each PAC-modulated population, giving Δ*d*′. This is done between each successive time-step to assess average Δ*d*′. Error bars are standard deviations. **c)** Examples of Poisson-distributed spike rates for the stimulus population *S* and each PAC-modulated population at a single time point. We compare 100 Poisson draws at (rate = 10), creating distribution *S* (light grey). *S* was then transformed into distributions *EI, E*, or *I* with each transform mimicking the appropriate neural arithmetic operation (dark grey; see *Methods*). As can be seen, the average firing rate for *S* is approximate equal to the defined rate of 10 spikes. In the EI/gain population (top), both the firing rate and variance of the downstream population are higher than the input drive, greatly improving the signal-to-noise ratio. In contrast, oscillatory E populations only show modest signal-to-noise improvements (middle), while I populations show small decreases (bottom).

### Information flow and signal detection

From the point of view of downstream neuron receiving action potentials, and from the point of an information theoretic framework, reliable information transmission requires a good estimate of conditional probabilities. That is, the system needs to determine if a change in firing rate is the result of a change to the underlying neural encoding, or due to noise-driven fluctuations (Figure 3**a**). When viewing neurons as transmission devices, ideal downstream neurons should only adjust their firing to changes in signal or encoding, and not noise. Likewise, our information theoretic analyses estimate both *p*(*r*), the probability of the system having rate *r* and *p*(*r|s*), the probability of *r* given some input stimulus, *s*. The more reliable these quantities are, the better our estimate of mutual information.

The relation between neural arithmetic and signal detection is clearly demonstrated by comparing Poisson distributed data for a single time point, *r*. We compare 100 Poisson draws at *r* = 2, creating distribution *S. S* is then transformed into distributions *EI, E*, or *I* by the appropriate neural arithmetic operation (see *Methods*). Figure 3**c** demonstrates that multiplicative gain leads to a strong increases in separation from *S*. In contrast additive and subtractive alterations to to *S*, akin to those present in models *E* and *I*, show no change or worsening separation, respectively (Figure 3**c**).

To quantify signal separation in our neural models we measured *d*′ (*Methods*) between each successive time-step, comparing the observed population rate for S to the three PAC models (Figure 3**b**). As expected, model *EI* showed consistent increases in *d*′, increasing as a function of stimulus drive (*r_s_*) and modulation strength (*g*). Model E had a slight increase, while I saw a slight decrease. When comparing trends in Δ*d*′ to Δ*ΜΙ* (*i.e.,* Figure 3**b** to 1**e**) there are two notable discrepancies. For model *EI*, the MI effect plateaus as a function of both *r_s_* and *g.* In contrast, *d*′ scales nearly linearly. This can be explained when considering the signal detection problem in relation to information theory.

We argue that the more reliable the estimate of *p*(*r*) and *p*(*r*|*s*), the better our ability to estimate MI. However this effect has a limit. Once the probability estimates of have become sufficiently reliable, further increases to *d*′ are of no benefit, leading Δ*ΜΙ* to plateau. Additionally, Δ*ΜΙ* decreases strongly for both *E* and *I*, but the effect on Δ*d*′ in these models is mild. However, recall that we compared the separation of only successive estimates in rate, but the oscillatory modulation spans multiple time-steps and introduces a very different pattern than that of naturalistic firing. Without the increase in signal separation to compensate, the modulatory oscillatory pattern corrupts our estimate of *p*(*r*) and *p*(*r*|*s*), giving the observed decrease in Δ*MI*.

## Discussion

In this modeling study we provide novel evidence that oscillatory modulation of a population of spiking neurons can actually enhance mutual information between an input signal and its neural encoding via improved neural signal detection. This enhancement requires that PAC act as a multiplicative gain modulator, a requirement synonymous with strong *and* balanced excitatory and inhibitory inputs [Chance et al., 2002, Vreeswijk and Sompolinsky, 1998, Brunel, 2000, Abbott and Chance, 2005, Atallah and Scanziani, 2009, Womelsdorf et al., 2014]. When there is an imbalance in either direction–such that PAC is driven by either excitatory or inhibitory inputs–information flow is obstructed. These results have striking implications for the role of neural oscillations in coordinating functional neural assemblies, and make a number of predictions regarding network dysfunction in neurological and psychiatric disease. Importantly, our model makes no *a priori* assumptions about the utility or function of oscillations; rather the results emerge from a purely analytic framework grounded in first principles of neuronal physiology. Our model unifies seemingly paradoxical results wherein PAC is related to successful cognitive function across multiple cognitive domains, but is also observed in neuropathological disorders including Parkinson’s disease [de Hemptinne et al., 2015], autism [Khan et al., 2013], and schizophrenia [Allen et al., 2011].

Prior empirical and modeling work suggests that gain control requires strong, balanced *EI* connections [Abbott and Chance, 2005, Lim and Goldman, 2013, Renart and Machens, 2014]. Interestingly, in healthy tissue there are momentary fluctuations in *EI* balance [Knoblich et al., 2010, Chance et al., 2002, Renart et al., 2007], however, under normal conditions homeostatic mechanisms keep these fluctuations from deviating too far. One of the implications of our results is that minor alterations in *EI* balance may allow for functional networks to switch between subassemblies, perhaps allowing for oscillatory multiplexing of population codes [Akam and Kullmann, 2014].

Within the E-I framework, oscillations are robust, with imbalances in either excitation or inhibition favoring their formation. Given their ubiquity, the question becomes how does neural entrainment by oscillations interact with imbalances in E-I? That is, not all oscillations are equal, and under *E* or *I* imbalance, *EI* multiplicative coupling between a stimulus and oscillatory populations is replaced with obstructive coupling.

This model provides a way for oscillations to be re-framed in terms of their informational utility. Thus, a number of neurological and psychiatric disorders that are associated with both *EI* imbalances at the microscale, and oscillatory abnormalities at the meso-and macro-scales, can now be assessed in terms of disruptions in the information flow within functional networks. Notably, a PAC network is itself an oscillator that can further entrain downstream neural populations. In this sense, *EI* imbalance may lead to a cascade of reduced neural information exchange. This is further exacerbated by the fact that E-I imbalance itself can favor the formation of oscillatory networks, resulting in the formation of sustained pathological information flow. This is strongly reminiscent of the pathologically strong oscillatory activity and *EI* imbalances seen in disorders such as Parkinson’s disease.

Additionally, our model successfully produces cross-frequency coupling between a low frequency phase and high gamma, a common method for assessing the functional role of oscillations. Importantly, however, cross-frequency coupling strength does not track changes to mutual information; more specifically, it does not track the temporal and spectral characteristics of gain modulation. This suggests a disassociation between spectral and functional changes.

It is notable that the system that yields the largest overall PAC is the inhibitory condition. However, despite this large PAC, there is no improvement in the mutual information between the input and the encoding population. This is likely due to the fact that the inhibitory system introduces a severe oscillation via periods of inhibition-induced silence, which comes at the cost of any possible encoding due to periods with no spiking. In contrast, in the excitatory condition there is an observed minor signal detection improvement, but relatively lower PAC. Here, it may be that relatively weaker inputs are needed to activate the encoding population, which, combined with weaker inhibitory modulation makes the population more susceptible to noise. This may yield a slight discrimination improvement, but at the cost of encoding accuracy.

It is common in the literature to relate increases in cross-frequency coupling to increases in communication efficiency. Our modeling suggests that this inference is not without peril. Our results (Figure 2) strongly suggest that, although oscillatory gain modulation has distinct spectral and temporal signatures, these features cannot be fully captured by narrowband cross-frequency coupling methods. Rather, the signatures for facilitative versus obstructive cross-frequency coupling may instead be best captured by studying interrelations of frequency bands in power spectra as part of a unified and temporally-sensitive system.

The time-courses of activity present under *EI, E*, and *I* conditions are distinct (Figure 1 and 4). Empirically, oscillatory “shape” (e.g., peakedness and asymmetry) vary by frequency, brain region, and behavioral state. However, there exists little modeling support to guide interpretation of these non-sinusoidal features in cognition, in terms of either biologically or functionally meaningful categories. Our results suggest that time-domain, non-sinusoidal features may be critical for understanding the relationship between PAC and information flow. In other words, the common Fourier-based spectral methods–which by definition have sinusoidal basis functions–may by their sinusoidal nature be masking our ability to detect biologically-relevant changes in neural information flow.

It is important to stress that any computational modeling approach has both strong benefits and limitations. The results presented herein would be difficult to experimentally assess given the biological difficulty in driving excitatory or inhibitory neurons with high specificity. In contrast, building this computational model allows us to explore multiple forms of biologically-plausible oscillations and their effects on information flow. However, any model is limited by its underlying assumptions. Here we conceive of spiking as a statistical (Poisson) process with simplified synaptic interactions. This comes at the cost of more complex neural dynamics such as changes to spike-timing induced by oscillatory activity or caused by the tuning to information flow that can follow from activity-dependent changes to synaptic strength (*i.e*., spike-time dependent plasticity). Nevertheless, our results offer a parsimonious explanation for a diversity of empirical PAC results in both cognition and disease. In conclusion, our model predicts that in a system of excitatory and inhibitory neurons, PAC will improve mutual information between an input stimulus and its neural encoding *only* when there is a excitation/inhibition balance. In contrast, when excitation or inhibition alone drives PAC, information flow is obstructed.

## References

Abbott, L. F. and Chance, F. (2005). Drivers and modulators from push-pull and balanced synaptic. Progress in Brain Research.

Ainsworth, M., Lee, S., Cunningham, M. O., Traub, R. D., Kopell, N. J., and Whittington, M. a. (2012). Rates and rhythms: a synergistic view of frequency and temporal coding in neuronal networks. Neuron, 75(4):572–83.

Akam, T. and Kullmann, D. M. (2014). Oscillatory multiplexing of population codes for selective communication in the mammalian brain. Nature reviews. Neuroscience, 15(2):111–22.

Allen, E. A., Liu, J., Kiehl, K. A., and Gelernter, J. (2011). Components of cross-frequency modulation in health and disease. Frontiers in systems {…}.

Atallah, B. V. and Scanziani, M. (2009). Instantaneous Modulation of Gamma Oscillation Frequency by Balancing Excitation with Inhibition. Neuron, 62(4):566–577.

Barbieri, F., Mazzoni, A., Logothetis, N. K., Panzeri, S., and Brunel, N. (2014). Stimulus Dependence of Local Field Potential Spectra: Experiment versus Theory. Journal of Neuroscience, 34(44):14589–14605.

Bays, P. M. (2014). Noise in neural populations accounts for errors in working memory. The Journal of neuroscience: the official journal of the Society for Neuroscience, 34(10):3632–45.

Börgers, C. and Kopell, N. (2003). Synchronization in networks of excitatory and inhibitory neurons with sparse, random connectivity. Neural computation, 15: 509–538.

Brunel, N. (2000). Dynamics of sparsely connected networks of excitatory and inhibitory spiking neurons. J of Computaional Neuroscience, 32–33:307–312.

Brunel, N. and Wang, X.-J. (2003). What determines the frequency of fast network oscillations with irregular neural discharges? I. Synaptic dynamics and excitation-inhibition balance. Journal of neurophysiology, 90(1):415–30.

Buehlmann, A. and Deco, G. (2010). Optimal information transfer in the cortex through synchronization. PLoS computational biology, 6(9).

Buschman, T. J., Denovellis, E. L., Diogo, C., Bullock, D., and Miller, E. K. (2012). Synchronous Oscillatory Neural Ensembles for Rules in the Prefrontal Cortex. Neuron, 76(4):838–846.

Buzsáki, G. (2010). Neural syntax: cell assemblies, synapsembles, and readers. Neuron, 68(3):362–85.

Buzsáki, G., Anastassiou, C. a., and Koch, C. (2012). The origin of extracellular fields and currents-EEG, ECoG, LFP and spikes. Nature reviews. Neuroscience, 13(6):407–20.

Buzsáki, G. C. N. K. S. U. O. R. O. U. A. T. F. Q. B. C. A. B. G. S. S. Q. B. C. D. U. E. -.-. K. S. U. M. S. B. G. Q. B. C. A. W. S. F. S. W. L.. (2006). Rhythms of the brain.

Canolty, R. T., Edwards, E., Dalal, S. S., Soltani, M., Nagarajan, S. S., Kirsch, H. E., Berger, M. S., Barbaro, N. M., and Knight, R. T. (2006). High gamma power is phase-locked to theta oscillations in human neocortex. Science (New York, N.Y.), 313(5793):1626–8.

Canolty, R. T. and Knight, R. T. (2010). The functional role of cross-frequency coupling. Trends in Cognitive Sciences, 14(11):506–515.

Cellulaire, L. D. N. (1992). Population Oscillations in the Ca3 Region of the in Vitro. pages 653–672.

Chance, F. S., Abbott, L. F., and Reyes, A. D. (2002). Gain modulation from background synaptic input. Neuron, 35(4):773–782.

Cooper, P. S., Wong, A. S. W., Fulham, W. R., Thienel, R., Mansfield, E., Michie, P. T., and Karayanidis, F. (2015). Theta frontoparietal connectivity associated with proactive and reactive cognitive control processes. NeuroImage, 108(C):354–363.

de Hemptinne, C., Ryapolova-Webb, E. S., Air, E. L., Garcia, P. a., Miller, K. J., Ojemann, J. G., Ostrem, J. L., Galifianakis, N. B., and Starr, P. a. (2013). Exaggerated phase-amplitude coupling in the primary motor cortex in Parkinson disease. Proceedings of the National Academy of Sciences of the United States of America, 110(12):4780–5.

de Hemptinne, C., Swann, N. C., Ostrem, J. L., Ryapolova-Webb, E. S., San Luciano, M., Galifianakis, N. B., and Starr, P. a. (2015). Therapeutic deep brain stimulation reduces cortical phase-amplitude coupling in Parkinson’s disease. Nature Neuroscience, 18(5).

Engel, a. K., Fries, P., and Singer, W. (2001). Dynamic predictions: oscillations and synchrony in top-down processing. Nature reviews. Neuroscience, 2(10):704–716.

Fell, J. and Axmacher, N. (2011). The role of phase synchronization in memory processes. Nature reviews. Neuroscience, 12(2):105–18.

Fries, P. (2005). A mechanism for cognitive dynamics: neuronal communication through neuronal coherence. Trends in Cognitive Sciences, 9(10):474–480.

Fries, P., Reynolds, J. H., Rorie, a. E., and Desimone, R. (2001). Modulation of oscillatory neuronal synchronization by selective visual attention. Science (New York, N.Y.), 291(5508):1560–1563.

Geisler, C., Brunel, N., and Wang, X.-J. (2005). Contributions of intrinsic membrane dynamics to fast network oscillations with irregular neuronal discharges. Journal of neurophysiology, 94(6):4344–61.

Hansel, D., Mato, G., and Meunier, C. (1995). Synchrony in excitatory neural networks. Neural computation, 7(2):307–337.

Heeger, D. J., Simoncelli, E. P., and Movshon, J. A. (1996). Computational models of cortical visual processing. Proceedings of the National Academy of Sciences of the United States of America.

Herrmann, C. S. and Demiralp, T. (2005). Human EEG gamma oscillations in neuropsychiatric disorders. Clinical Neurophysiology, 116(12):2719–2733.

Hölscher, C., Anwyl, R., and Rowan, M. J. (1997). Stimulation on the positive phase of hippocampal theta rhythm induces long-term potentiation that can Be depotentiated by stimulation on the negative phase in area CA1 in vivo. Journal of Neuroscience, 17(16):6470–6477.

Ince, R. a. a., Petersen, R. S., Swan, D. C., and Panzeri, S. (2009). Python for information theoretic analysis of neural data. Frontiers in neuroinformatics, 3(February):4.

Jensen, O., Idiart, M. A. P., and Lismanl, E. (1996). Physiologically Realistic Formation of Autoassociative Memory in Networks with Theta / Gamma Oscillations: Role of Fast NMDA Channels.

Jia, X., Tanabe, S., and Kohn, A. (2013). Gamma and the Coordination of Spiking Activity in Early Visual Cortex. Neuron, 77(4):762–774.

Khan, S., Gramfort, A., and Shetty, N. R. (2013). Local and long-range functional connectivity is reduced in concert in autism spectrum disorders. In Proceedings of the {…}.

Knoblich, U., Siegle, J. H., Pritchett, D. L., and Moore, C. I. (2010). What do we gain from gamma? Local dynamic gain modulation drives enhanced efficacy and efficiency of signal transmission. Frontiers in human neuroscience, 4(October):185.

Lim, S. and Goldman, M. S. (2013). Balanced cortical microcircuitry for maintaining information in working memory. Nature neuroscience, 16(9):1306–14.

Lisman, J. E. and Jensen, O. (2013). The *θ*-γ neural code. Neuron, 77(6):1002–16.

Litwin-Kumar, A. and Doiron, B. (2012). Slow dynamics and high variability in balanced cortical networks with clustered connections. Nature neuroscience, 15(11):1498–505.

Litwin-Kumar, A. and Doiron, B. (2014). Formation and maintenance of neuronal assemblies through synaptic plasticity. Nature communications, 5(May):5319.

Litwin-Kumar, A., Oswald, A.-M. M., Urban, N. N., and Doiron, B. (2011). Balanced synaptic input shapes the correlation between neural spike trains. PLoS computational biology, 7(12):e1002305.

MacLeod, K. and Laurent, G. (1996). Distinct mechanisms for synchronization and temporal patterning of odor-encoding neural assemblies. Science, 274: 976–979.

Mazurek, M. E. and Shadlen, M. N. (2002). Limits to the temporal fidelity of cortical spike rate signals. Nature neuroscience, 5(5):463–71.

Mazzoni, A., Brunel, N., Cavallari, S., Logothetis, N. K., and Panzeri, S. (2011). Cortical dynamics during naturalistic sensory stimulations: Experiments and models. Journal of Physiology Paris, 105(1–3):2–15.

Murphy, B. K. and Miller, K. D. (2009). Balanced Amplification: A New Mechanism of Selective Amplification of Neural Activity Patterns. Neuron, 61(4):635–648.

N. E. Huang, Z. Shen, S. Long, M. Wu, H. Shih, Q. Zheng, N.-C. Yen, C. Tung, and H. Liu (1998). The empirical mode decomposition and the Hilbert spectrum for nonlinear and non-stationary time series analysis. Proceedings of the Royal Society of London. Series A: Mathematical, Physical and Engineering Sciences, 454(1971):903–995.

Ostojic, S. (2014). Two types of asynchronous activity in networks of excitatory and inhibitory spiking neurons. Nature neuroscience, 17(4):594–600.

Özkurt, T. E. and Schnitzler, A. (2011). A critical note on the definition of phase-amplitude cross-frequency coupling. Journal of Neuroscience Methods, 201(2):438–443.

Panzeri, S. and Treves, A. (1996). Analytical estimates of limited sampling biases in different information measures. Network: Computation in Neural Systems, 7(1):87–107.

Renart, A. and Machens, C. K. (2014). Variability in neural activity and behavior. Current Opinion in Neurobiology, 25: 211–220.

Renart, A., Moreno-Bote, R., Wang, X.-J., and Parga, N. (2007). Mean-driven and fluctuation-driven persistent activity in recurrent networks. Neural computation, 19(1):1–46.

Reynolds, J. H. and Heeger, D. J. (2009). The Normalization Model of Attention.

Saalmann, Y. B., Pinsk, M. a., Wang, L., Li, X., and Kastner, S. (2012). The Pulvinar Regulates Information Transmission Between Cortical Areas Based on Attention Demands. Science, 337(6095):753–756.

Schaefer, A. T., Angelo, K., Spors, H., and Margrie, T. W. (2006). Neuronal oscillations enhance stimulus discrimination by ensuring action potential precision. PLoS Biology, 4(6):1010–1024.

Siegel, M., Buschman, T. J., and Miller, E. K. (2015). Cortical information flow during flexible sensorimotor decisions. Science, 34: 1–5.

Silver, R. A. (2010). Neuronal arithmetic. Nature reviews. Neuroscience, 11(7):474–89.

Sohal, V. S., Zhang, F., Yizhar, O., and Deisseroth, K. (2009). Parvalbumin neurons and gamma rhythms enhance cortical circuit performance. Nature, 459(7247):698–702.

Spaak, E., de Lange, F. P., and Jensen, O. (2014). Local Entrainment of Alpha Oscillations by Visual Stimuli Causes Cyclic Modulation of Perception. Journal of Neuroscience, 34(10):3536–3544.

Szczepanski, S. M., Crone, N. E., Kuperman, R. A., Auguste, K. I., Parvizi, J., and Knight, R. T. (2014). Dynamic Changes in Phase-Amplitude Coupling Facilitate Spatial Attention Control in Fronto-Parietal Cortex. PLoS Biology, 12(8):e1001936— -14.

Tiesinga, P. and Sejnowski, T. J. (2009). Cortical enlightenment: are attentional gamma oscillations driven by ING or PING? Neuron, 63(6):727–32.

Tort, A. B. L., Komorowski, R., Eichenbaum, H., and Kopell, N. (2010). Measuring phase-amplitude coupling between neuronal oscillations of different frequencies. Journal of neurophysiology, 104(2):1195–210.

Tort, A. B. L., Komorowski, R. W., Manns, J. R., Kopell, N. J., and Eichenbaum, H. (2009). Theta-gamma coupling increases during the learning of item-context associations. Proceedings of the National Academy of Sciences of the United States of America, 106(49):20942–20947.

Uhlhaas, P. J. and Singer, W. (2006). Neural Synchrony in Brain Disorders: Relevance for Cognitive Dysfunctions and Pathophysiology. Neuron, 52(1):155–168.

von Nicolai, C., Engler, G., Sharott, A., Engel, A. K., Moll, C. K., and Siegel, M. (2014). Corticostriatal Coordination through Coherent Phase-Amplitude Coupling. The Journal of neuroscience: the official journal of the Society for Neuroscience, 34(17):5938–48.

Voytek, B., D’Esposito, M., Crone, N., and Knight, R. T. (2013). A method for event-related phase/amplitude coupling. NeuroImage, 64: 416–424.

Voytek, B., Kayser, A. S., Badre, D., Fegen, D., Chang, E. F., Crone, N. E., Parvizi, J., Knight, R. T., and D’Esposito, M. (2015). Oscillatory dynamics coordinating human frontal networks in support of goal maintenance. Nature Neuroscience.

Voytek, B. and Knight, R. T. (2010). Prefrontal cortex and basal ganglia contributions to visual working memory. Proceedings of the National Academy of Sciences of the United States of America, 107(42):18167–72.

Voytek, B. and Knight, R. T. (2015). Dynamic network communication as a unifying neural basis for cognition, development, aging, and disease. Biological Psychiatry, X(X).

Vreeswijk, C. V. and Sompolinsky, H. (1998). Chaotic Balanced State in a Model of Cortical Circuits. Neural computation, 1371: 1321–1371.

Wang, X.-J. (2010). Neurophysiological and computational principles of cortical rhythms in cognition. Physiological reviews, 90(3):1195–268.

Wang, X.-J. and Buzsaki, G. (1996). Gamma Oscillation by Synaptic Inhibition in a Hippocampal Interneuronal Network Model. J. Neurosci., 16(20):6402–6413.

Womelsdorf, T., Schoffelen, J.-M., Oostenveld, R., Singer, W., Desimone, R., Engel, A. K., and Fries, P. (2007a). Modulation of neuronal interactions through neuronal synchronization. Science (New York, N.Y.), 316(June):1609–1612.

Womelsdorf, T., Schoffelen, J.-M., Oostenveld, R., Singer, W., Desimone, R., Engel, A. K., and Fries, P. (2007b). Modulation of neuronal interactions through neuronal synchronization. Science (New York, N.Y.), 316(June):1609–1612.

Womelsdorf, T., Valiante, T. a., Sahin, N. T., Miller, K. J., and Tiesinga, P. (2014). Dynamic circuit motifs underlying rhythmic gain control, gating and integration. Nature Neuroscience, 17(8):1031–1039.

